# Protein noise and distribution in a two-stage gene-expression model extended by an mRNA inactivation loop

**DOI:** 10.1101/2021.04.22.440897

**Authors:** Candan Çelik, Pavol Bokes, Abhyudai Singh

## Abstract

Chemical reaction networks involving molecular species at low copy numbers lead to stochasticity in protein levels in gene expression at the single-cell level. Mathematical modelling of this stochastic phenomenon enables us to elucidate the underlying molecular mechanisms quantitatively. Here we present a two-stage stochastic gene expression model that extends the standard model by an mRNA inactivation loop. The extended model exhibits smaller protein noise than the original two-stage model. Interestingly, the fractional reduction of noise is a non-monotonous function of protein stability, and can be substantial especially if the inactivated mRNA is stable. We complement the noise study by an extensive mathematical analysis of the joint steady-state distribution of active and inactive mRNA and protein species. We determine its generating function and derive a recursive formula for the protein distribution. The results of the analytical formula are cross-validated by kinetic Monte-Carlo simulation.

## 1 Introduction

As many other biochemical mechanisms, gene expression in which protein synthesis occurs is inherently stochastic due to random fluctuations in the copy number of gene products, e.g. proteins [7]. From the viewpoint of biochemical reactions, in simplest formulations, gene expression consists of two main steps: transcription and translation. While RNA polymerase enzymes produce mRNA molecules in the former, protein synthesis takes place by ribosomes in the latter, each reaction corresponding to the production and decay of relevant species. Additionally, the two-stage model can be extended by the regulation of transcription factors, which affect gene expression by modulating the binding rate of RNA polymerase [3].

Over the last decades, the two-stage model of gene expression has been extensively studied to understand how the stochastic phenomenon in cellular processes takes place [14, 17, 18]. Specifically, quantifying the number of species in terms of probability distributions has become an interesting and challenging endeavour due to the subtleties involved in finding a solution to the underlying problem. On the other hand, the fluctuations in mRNA and protein levels are considered as a major source of noise, leading to cell-to-cell variability in gene regulatory networks [12, 15, 16]. The noise emerges from different sources, namely *intrinsic* and *extrinsic* noise [23, 25]; yet, structural elements such as stem-loops can also contribute to noise by binding to an untranslated region of mRNA [6]. The untranslated regions of mRNAs often contain these stem-loops that can reversibly change configurations making individual mRNAs translationally active/inactive.

Numerous modelling approaches have been proposed that are based on deterministic and stochastic frameworks, and recently also hybrid ones as a combination of the preceding two [5, 10, 21]. Only a few of those provide an explicit solution to the two-stage gene-expression model [4, 18]; most of the studies are based on Monte Carlo simulations, which are usually computationally expensive.

As a generalisation of the two-state model, some studies in the literature consider a set of multiple gene states and investigate the dynamics of stochastic transitions among these states [11, 26]. Nevertheless, to the best of our knowledge, none of these studies takes an mRNA inactivation into account. Here we extend the two-stage model by an MRNA inactivation loop, by which we mean that after transcription species can switch between active and inactive states. In other words, there exists a pair of reversible chemical reactions occurring at constant rates by turning active mRNA species into inactive ones, and vice versa. Subsequently, the active mRNA is translated, while the inactive mRNA stays dormant. The schematic of reactions describing the model is given in (1). Here we thereafter refer to the aforementioned model as *the extended model*.

This paper is organised as follows. In Section 2, the stationary means of active mRNA, inactive mRNA, and protein are obtained from a deterministic formulation the model; the master equation of the stochastic model is formulated, and transformed into a partial differential equation for the generating function. In Section 3, the partial differential equation is transformed into one for the factorial cumulant generation function and a power series solution is found; recursive expressions for the coefficients — the factorial cumulants of the three molecular species — are thereby provided. In Section 4, the protein Fano factor is expressed in terms of the first two factorial cumulants, and the noise-reduction effect of the mRNA inactivation loop is analysed. The generating function of the stationary distribution of active mRNA, inactive mRNA and protein amounts is represented in the special-function form in Section 5. The marginal protein and active and inactive mRNA distributions are derived in Section 6. The paper is concluded in Section 7.

## 2 Model formulation

The extended model involves three species, mRNA, inactive mRNA (imRNA for short), and protein, and consists of the reactions

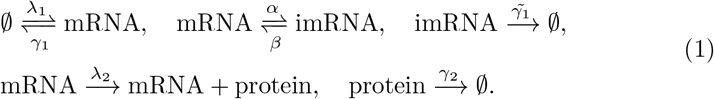

The reactions in (1) correspond to mRNA transcription and decay, mRNA activation and inactivation, inactive mRNA decay, protein translation, and protein decay, respectively.

Due to the linearity of kinetics in (1), the mean levels of the mRNA (*m*), inactive mRNA 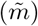 and protein (*n*) exactly satisfy the system of deterministic rate equations

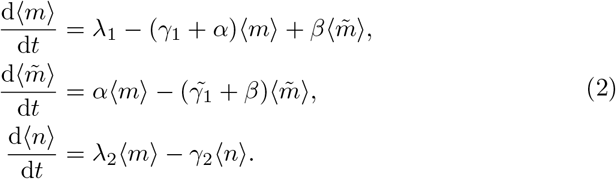

Setting time derivatives in (2) to zero, and solving the resulting algebraic system, the stationary means are obtained as

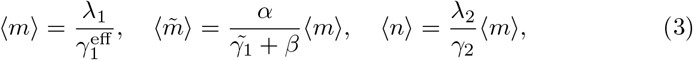

for the mRNA, inactive mRNA, and protein respectively, where

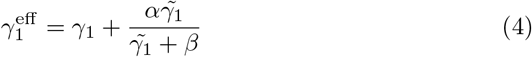

denotes the effective rate of mRNA decay. Owing to the linearity of reaction rates, one can find a closed system of differential equations not only for means, but also for higher-order moments [19, 22]; however these equations are typically less revealing than the mean dynamics. Here we take a different approach and quantify the protein noise as a by-product of a generating-function analysis in Section 4.

The probability 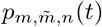 of having *m* mRNA, 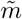 inactive mRNA, and *n* protein molecules at time *t* satisfies the chemical master equation

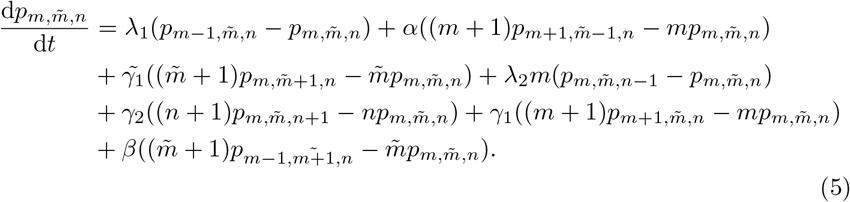

Equating the left-hand side of (5) to zero yields the steady-state master equation

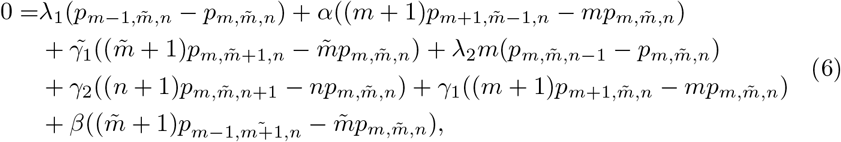

We additionally require that the normalising condition

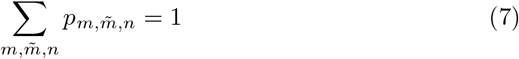

hold.

We aim to find the moments of the probability distribution 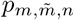 by using the generating function approach [8]. In order to solve (6)–(7), we employ the probability generating function

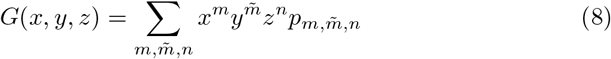

for the probability distribution 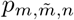. Multiplying (6) by the factor 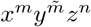 and summing over *m*, 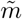 and *n* yields

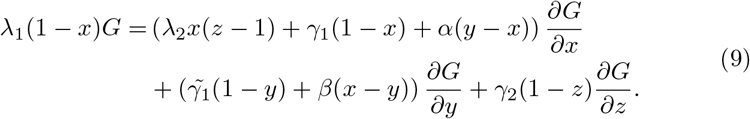

Equation (9) is subject to

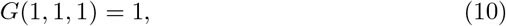

which is implied by the normalisation condition (7).

## 3 Factorial cumulant generating function

In order to find a particular solution to (9)–(10), we change the variables according to

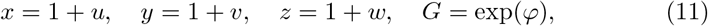

and obtain that the factorial cumulant generating function [9] *φ* = *φ*(*u, v, w*) is a solution of the inhomogeneous linear partial differential equation (PDE),

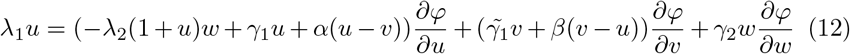

subject to

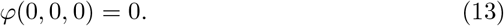

In order to solve (12)–(13) we shall employ the ansatz

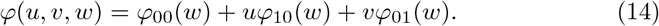

We immediately obtain the partial derivatives

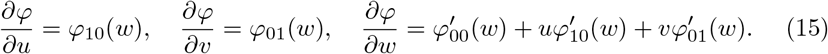

Inserting (15) into (12) and rearranging the terms yields an inhomogeneous system of ODEs

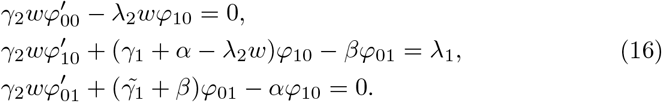

Let us assume that the functions *φ*_00_, *φ*_10_, and *φ*_01_ are of the power series form, i.e.,

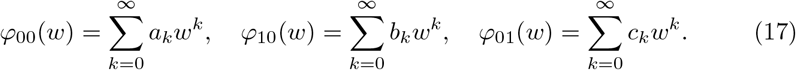

The coefficients *a_k_*, *b_k_*, and *c_k_* give the factorial cumulants of the joint molecular distribution [9]. Note that *a*_0_ = 0 follows immediately from the normalisation condition (13). Evaluating the derivatives in (17) and substituting into (16), we obtain the following recurrence equations:

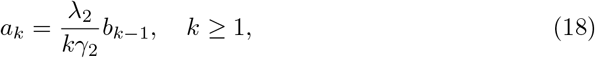

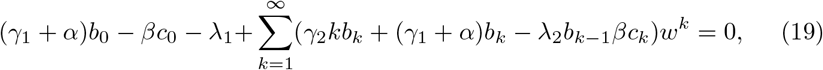

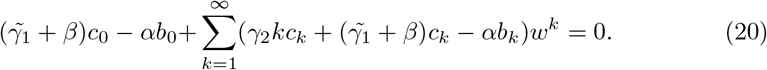

Since we consider (17) as a solution to (12) then all the coefficients in (19)–(20) must be zero. Thus, we get

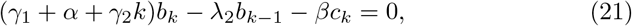

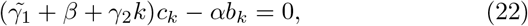

for *b_k_* and *c_k_*. Solving the algebraic system (21)–(22) in *b_k_*, *k ≥* 1, yields

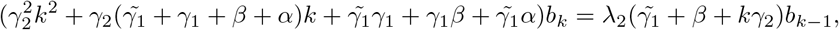

i.e.

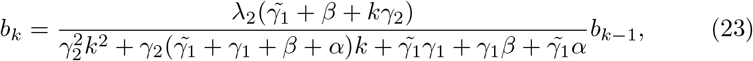

where the zeroth term of the sequence *b_k_* is obtained, by equating the terms out of the sums in (19) and (20) to zero, as

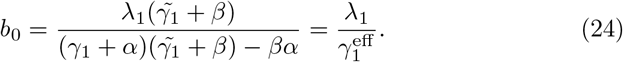

Equation (24) thus rederives the stationary mRNA mean (3) by means of factorial cumulant analysis; similarly, *c*_0_ and *a*_1_ can be identified as the stationary imRNA and protein means. Thus, the sequence *b_k_* can be calculated iteratively from (23) starting from the initial condition (24). Having calculated *b_k_*, the sequence *a_k_* and *c_k_* can be evaluated via (18) and (22). In Section 5, we will utilise these formulas to obtain a special-function representation of the generating function. Before doing that, we show that the first two terms of these sequences determine protein variability.

## 4 Protein variability

As outlined in the previous section, the first-order cumulants *b*_0_, *c*_0_, and *a*_1_ (*a*_0_ = 0 by normalisation condition), coincide with the stationary mRNA, imRNA, and protein mean values. In this section, we use the second-order cumulants to describe the stationary noise in our model. The noise in mRNA and imRNA is Poissonian (see Section 6 for details) and therefore uninteresting: we focus on the protein noise.

This section is divided into two parts: the first expresses the Fano factor in terms of the first and second order cumulants (and is independent of the specifics of the current model); the second part uses the formula to analyse the noise reduction effect of the inactivation loop.

### Expressing the Fano factor in terms of the cumulants

The generating function is expanded by the Taylor formula as

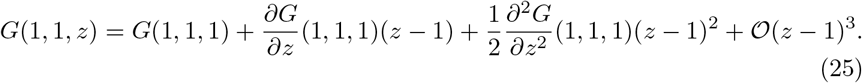

Differentiating (8) with respect to *z* and setting (*x, y, z*) = (1, 1, 1) links the derivatives of the generating function to the factorial moments:

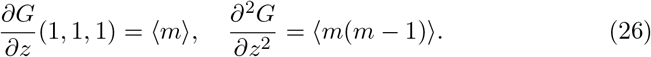

Inserting (10) and (26) into (25), we have

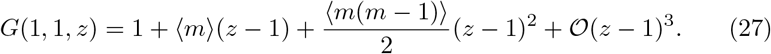

On the other hand, (11), (14), and (17) imply

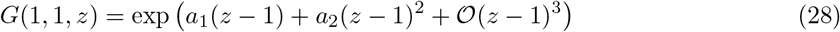

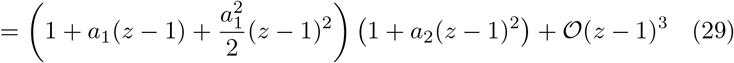

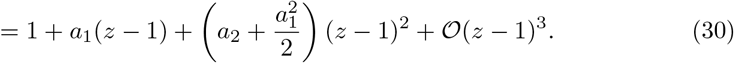

Comparing (27) and (28) gives

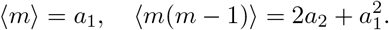

The Fano factor,

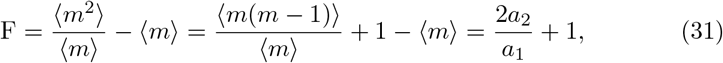

is thus expressed in terms of the first two factorial cumulants *a*_1_ and *a*_2_.

### Noise reduction by mRNA inactivation loop

Substituting (18) and (23) into (31) and simplifying gives

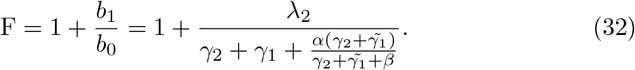

Formula (32) gives the steady-state protein Fano factor as function of the model parameters (degradation rate constants *γ*_1_, 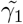, *γ*_2_ of active/inactive mRNA and protein; inactivation/activation rate constants *α, β*; translation rate constant *λ*_2_).

In order to compare the protein noise in the current model to that exhibited by the classical two-stage model (without the inactivation–activation loop) we define the baseline Fano factor as

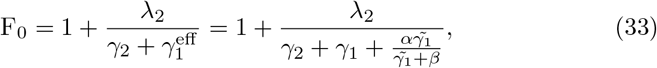

which can be obtained from (32) by first setting *α* = 0 (no inactivation) and then replacing the mRNA decay rate *γ*_1_ by its effective value (4). Adjusting the mRNA decay rate maintains the same species means in the baseline model like in the full model extended by the inactivation loop.

The protein variability formulae (32) and (33) can equivalently be expressed in terms of the squared coefficient of variation [13, 20] CV^2^ = F*/⟨n⟩* and 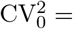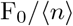. Combining (3) and (32)–(33), we find

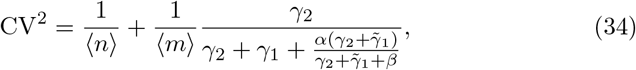

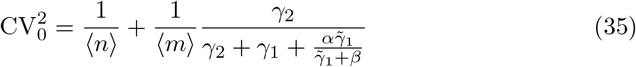

for the protein coefficient of variation and its baseline value (no activation loop).

Comparing (34) to (35), we see that 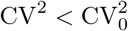, allowing us to conclude that the inclusion of the mRNA inactivation loop decreases protein noise. However, the two coefficients will be very close in many parameter regimes; the necessary conditions for observing a significant difference are given by

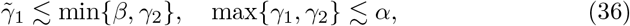

where by “≲” we mean smaller than or of similar magnitude. Thus, in order to obtain significant reduction of noise, we require that an individual active mRNA molecule be more likely to be inactivated than degraded, and that an individual inactive mRNA molecule be more likely to be activated than degraded. Additionally, we require that inactive mRNA be more stable than protein (which is possible if inactivation protects the mRNA from decay).

One particular consequence of the necessary conditions (36) is that the fraction protein noise reduction, 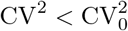, is a non-monotonous function of protein stability: it tends to one for highly unstable or highly stable proteins, and is less than one for proteins of optimal stability (cf. Figure 1). The optimal value of protein stability critically depends on the rate constant *β* of mRNA activation. In case of fast mRNA activation, the optimum noise reduction is achieved by unstable proteins (less stable than mRNA; Figure 1, left panel. In case of slow mRNA activation, the optimum can be achieved by stable proteins (Figure 1, right panel). However, slow activation (*β* 1) imposes, via (36), a stringent condition on the stability of inactivated mRNA. Indeed, the right panel of Figure 1 demonstrates that there is hardly any reduction of noise if the inactive mRNA is unstable.

**Fig. 1:**
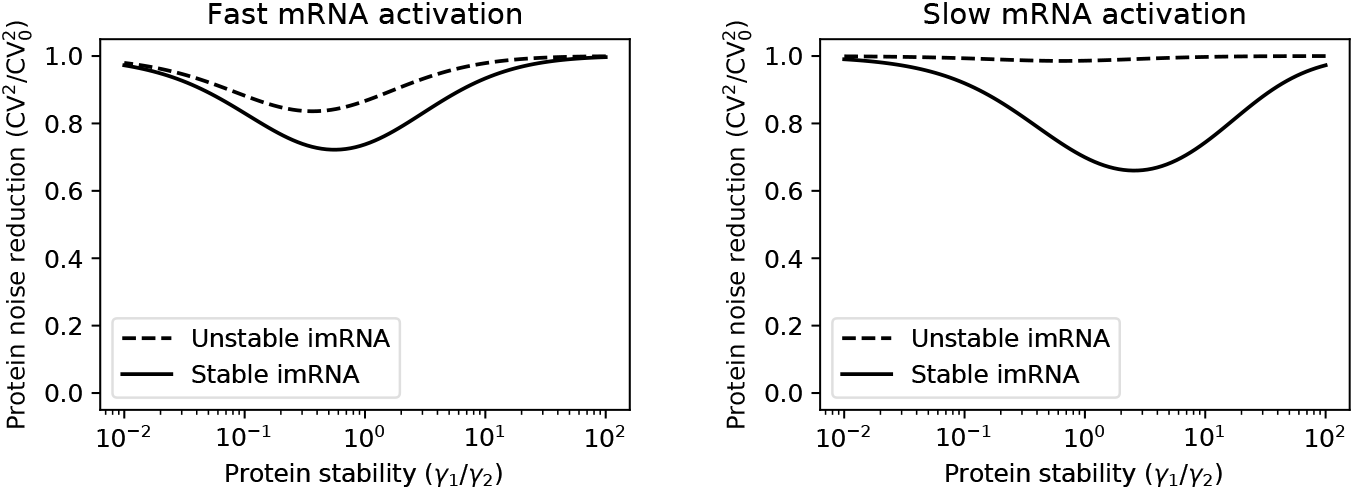
Fractional protein noise reduction by the mRNA inactivation loop as function of protein stability. The ordinate gives the protein noise (the squared coefficient of variation) in the two-stage model extended by the mRNA inactivation loop relative to the protein noise in a baseline two-stage model without the mRNA inactivation loop (adjusting the mRNA decay rate to obtain the same species means). The protein mean is set to *⟨n⟩* = 500; the mRNA mean is *⟨m⟩* = 10; the imRNA decay rate is either the same as that of active mRNA 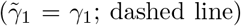 or set to zero 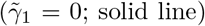. The inactivation and activation rates are *α* = 3, *β* = 3 (left panel) or *α* = 1, *β* = 0.1 (right panel); we thereby set *γ*1 = 1 without loss of generality.

In the next section, we go beyond the mean and noise statistics (the first and second order factorial cumulants), using the higher order cumulants to find a special-function representation of the generating function of the joint distribution of mRNA, imRNA, and protein copy numbers.

## 5 Special-function representation

Factorising the second-order polynomial in *k* in the denominator of (23) gives

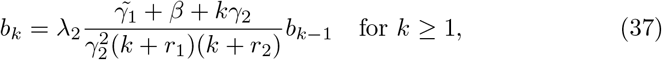

where

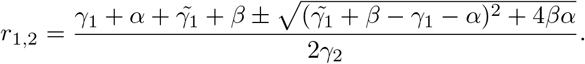

Note that the sequence *b_k_* in (37) can be rewritten as

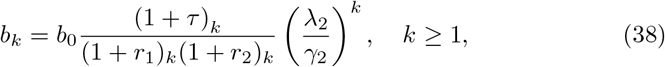

where we set 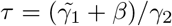 for the sake of simplicity and the polynomial

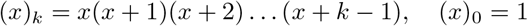

represents the rising factorial, also called the Pochammer symbol.

We next find the remaining sequences *a_k_* and *c_k_*. Inserting (38) into (18) gives

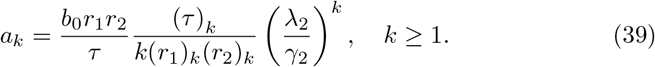

Similarly, substituting (38) into (22) yields

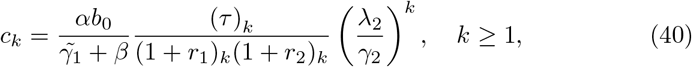

where 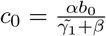, which can be obtained by combining (20) and (24).

Having found the sequences in (17), we next return to the original variables in (11) to obtain the generating function of the stationary distribution of active mRNA, inactive mRNA, and protein amounts, which is given by

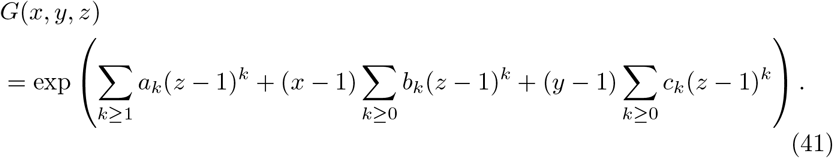

Equation (41) can be rewritten as

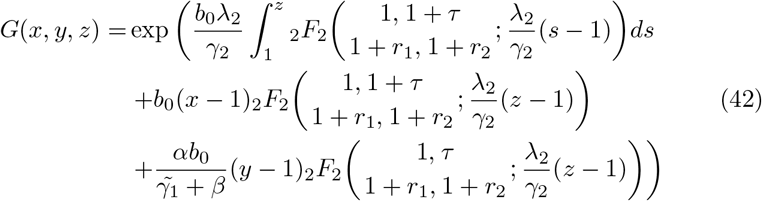

in terms of the generalised hypergeometric functions defined by [2]

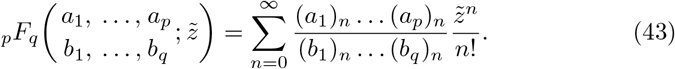

Equation (41) provides the sought-after special function representation of the joint generating function. In the following section, we focus on specific one-dimensional sections of the joint generating function that give the generating functions of the three marginal distributions.

## 6 Marginal distributions

In this section, we use the analytic formula (42) for the generating function to determine the marginal active and inactive mRNA, and protein distributions. To do so, we first set *y* = *z* = 1 in (42) and obtain

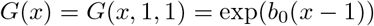

for the marginal active mRNA distribution. Similarly, setting *x* = *z* = 1 in (42) yields the marginal inactive mRNA distribution

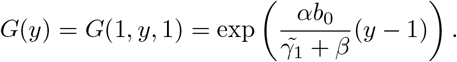

Finally, we set *x* = *y* = 1 in (42) and get the marginal protein generating function *G*(*z*) as

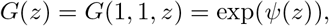

where *φ* is given by

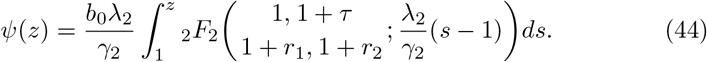

In order to obtain the marginal protein distribution, we exploit its generating function

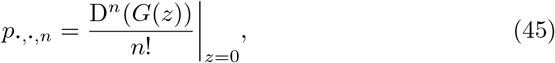

where D stands for the differential operator *d/dz* and 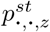 gives the probability of having *z* protein molecules and any number of active and inactive amount of mRNA. The first derivative of the composite function *G*(*z*) in (45) is obtained by chain rule as

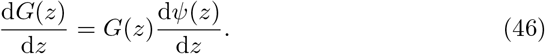

For the *n*-th derivative, we evaluate the (*n*1)th derivative of (46) according to the Leibniz rule, thus we have

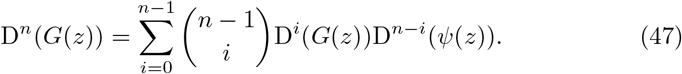

Next, we determine the *r*th–*r* is an arbitrary positive integer–derivative of the function *ψ*(*z*), which is given by

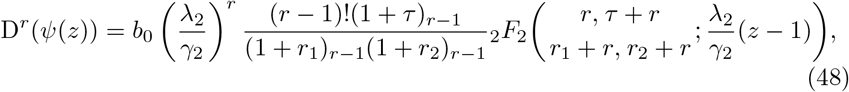

in which we used the formula

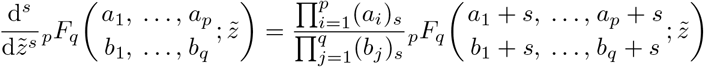

for the *s*-th derivative of the generalised hypergeometric function *_p_F_q_*. Inserting the derivatives in (48) into (47), taking *z* = 0, and rearranging the resulting equation according to (45) gives the formula for the marginal protein probabilities

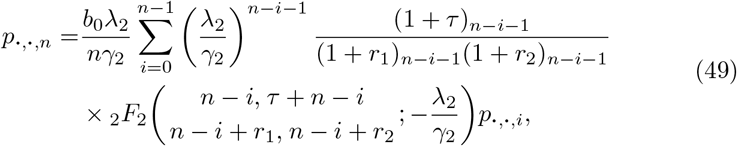

where the first term of the series is given by

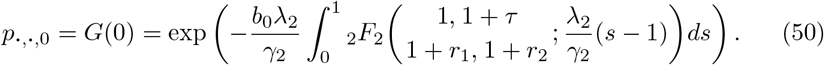

In order to calculate and compare the marginal protein probabilities (49) with those obtained by stochastic simulations based on Gillespie’s algorithm, we implement the recursive formula (49) in a high-level programming language, Python, together with using its numerical computing library NumPy and plotting library Matplotlib. The probabilities in (49) are calculated iteratively starting from its first term given by (50) up to *n* = 50. In Figure 2, the right panel compares the theoretical probability distribution (49) (blue bars) with the one obtained using stochastic simulations (solid line) at the timepoint *t* = 100, while the left panel shows the same comparison but on a logarithmic scale. The number of Gillespie iterations was set to 105 in the Python package GillesPy2 [1]. The initial number of active and inactive mRNA and protein was set to 5. A Python routine `mpmath.hyp2f2` used to calculate the generalised hypergeometric function _2_*F* in (49)–(50).

**Fig. 2:**
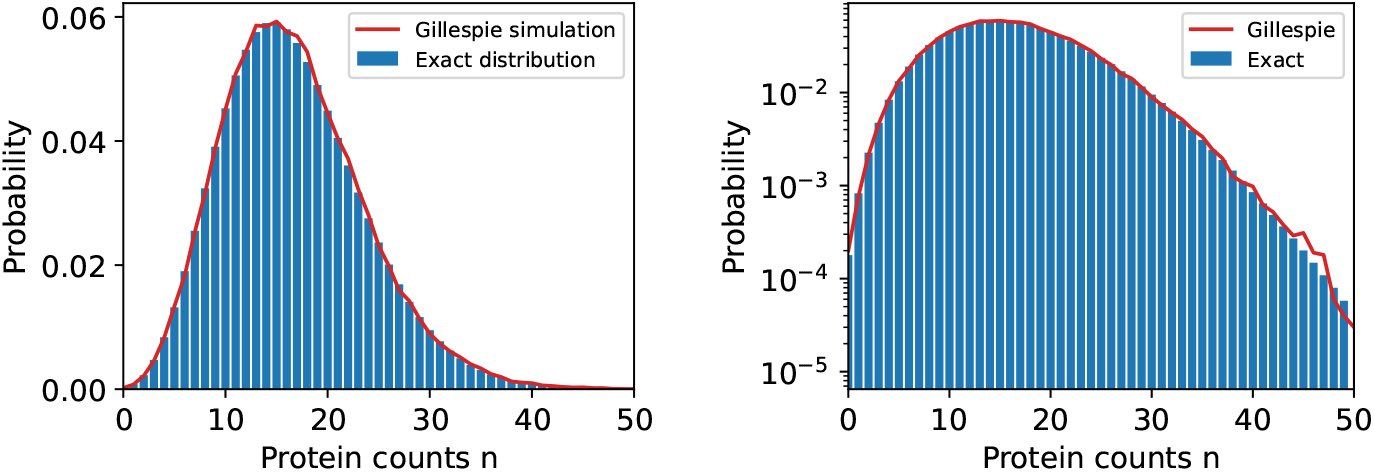
*Left:* Comparison of the probability mass function (49) of the marginal protein distribution and the probability calculated by Gillespie’s stochastic simulation algorithm (the solid line). *Right:* A logarithmic scale plot of the probability, out of 105 repeats, obtained by the two approaches. *Parameter values:* The kinetic parameters are: 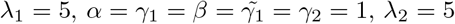.

## 7 Conclusion

In this paper, we analysed a formulation of the two-stage model for gene expression that extends the classical version [4, 24] by an mRNA inactivation loop. The principal results of our analysis are the characterisation of the mean and noise behaviour, as well as the underlying probability distribution. The principal tool is the factorial cumulant generating function and the factorial cumulant expansion.

The incorporation of the mRNA inactivation loop into the classical two-stage model for gene expression reduces the protein noise. However, in order for the reduction be substantial, several restrictions on the parameter rates have to be in place. In particular, the protein cannot be too stable or unstable, but its stability has to be optimally chosen. The resulting optimal value of protein stability is typically unrealistically low (lower than mRNA stability, in particular). In order to obtain an optimal stability that is greater than mRNA stability, one has to assume that inactivation protects the mRNA from degradation and activation is slow. Thus, our noise analysis points towards a potential role of the mRNA inactivation loop in gene expression noise control; at the same time, it delineates the limits of its application.

In addition to the noise analysis, we provide a comprehensive classification of the underlying probability distributions. Unsurprisingly, the distributions of the active/inactive mRNA are Poissonian. On the other hand, the protein distribution is highly non-trivial, and is characterised in terms of the generalised hypergeometric series. The characterisation is used to derive a recursive expression for the protein probability mass function. The recursive formula is found to be consistent with kinetic Monte-Carlo simulation (by means of the Gillespie direct method).

In summary, the paper provides a systematic mathematical analysis of an mRNA–protein model for gene expression extended by an inactive mRNA species, and hints at possible functional roles of mRNA inactivation loop in the control of low copy number gene-expression noise.

